# Chromosome-level genome assembly of a butterflyfish, *Chelmon rostratus*

**DOI:** 10.1101/719187

**Authors:** Xiaoyun Huang, Yue Song, Suyu Zhang, A Yunga, Mengqi Zhang, Yue Chang, He Zhang, Chang Li, Yong Zhao, Meiru Liu, Inge Seim, Guangyi Fan, Xin Liu, Shanshan Liu

**Author notes:** These authors contributed equally to this work (Xiaoyun Huang, Yue Song, Suyu Zhang, A Yunga). These authors jointly supervised this work (Xin Liu, Shanshan Liu). Corresponding to (S.L.) and (X.L.).

## Abstract

*Chelmon rostratus* (Teleostei, Perciformes, Chaetodontidae) is a copperband butterflyfish. As an ornamental fish, the genome information for this species might help understanding the genome evolution of Chaetodontidae and adaptation/evolution of coral reef fish.

In this study, using the stLFR co-Barcode reads data, we assembled a genome of 638.70 Mb in size with contig and scaffold N50 sizes of 294.41 kb and 2.61 Mb, respectively. 94.40% of scaffold sequences were assigned to 24 chromosomes using Hi-C data and BUSCO analysis showed that 97.3% (2,579) of core genes were found in our assembly. Up to 21.47 % of the genome was found to be repetitive sequences and 21,375 protein-coding genes were annotated. Among these annotated protein-coding genes, 20,163 (94.33%) proteins were assigned with possible functions.

As the first genome for Chaetodontidae family, the information of these data helpfully to improve the essential to the further understanding and exploration of marine ecological environment symbiosis with coral and the genomic innovations and molecular mechanisms contributing to its unique morphology and physiological features.

## Background & Summary

*Chelmon rostratus* (Teleostei, Perciformes, Chaetodontidae) lives in the western pacific, from southern Japan and Taiwan throughout the Coral Triangle to the Solomon Islands and the northern coast of Australia ^1^. In the natural environment, it lives in depths from 1- 25 meters underwater, inhabiting coastal and inner reefs and often in turbid water ^2^. It usually has an appealing appearance of four yellowish or yellowish-orange vertical bands with black border on the silvery white body, and a false eyespot. Thus, it is one of the protagonists in tropical marine aquarium fish ^3^. Despite as a common coral reef fish, there were limited previous researches in this species. Previous researches were limited to its behavior features such as the prey capture kinematics ^4^. More recently, its complete mitochondrial genome was reported ^5^, revealing its evolutionary position. However, there was no particular study providing the whole genomic information.

Here, we reported the first whole genome sequencing data for this species, resulting in a chromosome level genome assembly and annotation, followed by the evolutionary analysis. The data and analysis provided here, can benefit future basic studies and conservation efforts of this species.

## Methods

### Sample collection, library construction and sequencing

We captured an adult *Chelmon rostratus* (Figure 1) in the sea area of Qingdao, China, and muscle tissue was stored in liquid nitrogen. Then, high quality DNA was extracted using a modified DNA extraction for vertebrate tissues protocol ^6^ from the muscle tissue. The exacted DNA was fragmented and MGIEasy stLFR Library preparation kit (PN : 1000005622) was used to construct single tube Long Fragment Read (stLFR) library, following the stLFR protocol ^7^. For Hi-C library sequencing, about 1g living muscle tissue was used to DNA extraction and library contraction, according to Wang’s method ^7^. Sequencing was conducted on a BGISEQ-500 sequencer, generating 286.82Gb raw data (including 134.18Gb raw Hi-C data) (Supplementary Table 1). Data filtering was then carried out using SOAPnuke software (version 1.5) ^8^ with the default parameters, thus low-quality reads (more than 40% bases with quality score lower than 8), PCR duplications, adaptors and reads with high proportion (higher than 10%) of ambiguous bases (Ns) were filtered. After data filtering, 192.71 Gb ‘clean data’ (including 133.17Gb ‘clean Hi-C data’) was obtained for the further assembly (Supplementary Table 1).

**Figure 1.**
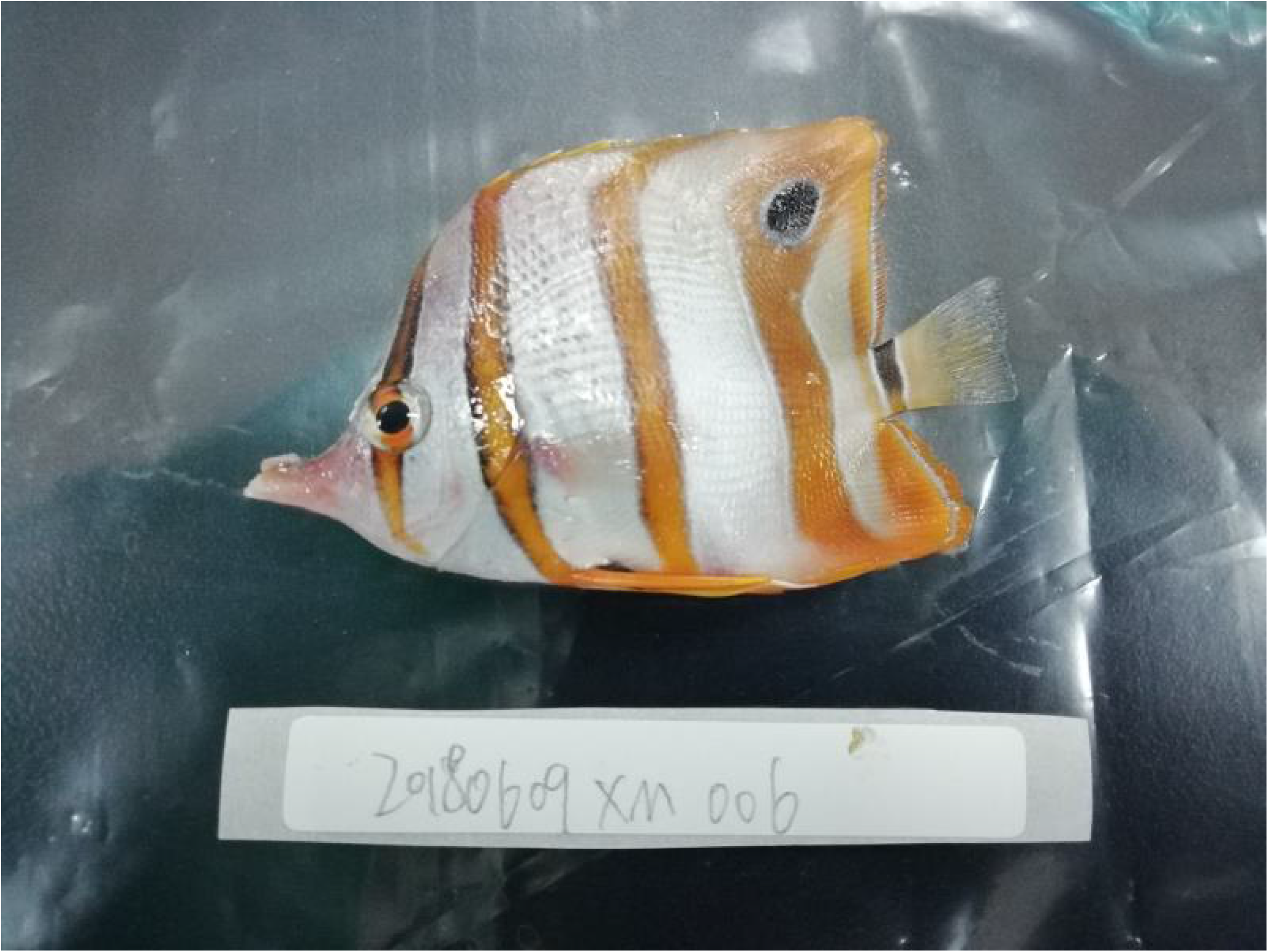
Photograph of *Chelmon rostratus*.

### Genome features revealed by *k*-mer analysis

In order to understand better about its genome features, we applied *k*-mer frequency distribution analysis to estimate its genome size and genome complexity (heterozygosity or/and repetition) ^9^. We randomly selected 59.54 Gb (~90X) clean reads and carried out a 17-mer analysis using KMERFREQ_AR (version 2.0.4) ^10^. The genome size was estimated to be ~711.39 Mb (Supplementary Table 2). Observing the distribution of 17-mer frequency (Supplementary Figure 1), we anticipated the genome to be a diploid species with slight heterozygosity proportion. In order to get the percentage of heterozygosity, GCE software ^9^ was used to calculation with 59.54 Gb clean reads and resulted that the percentage of heterozygosity was 0.72%.

### Genome assembly and annotation

We assembled the genome using Supernova (version 2.1) ^11^ with default parameters for 55 Gb clean stLFR data. After that, we used Gapcloser software ^10^ to fill the gaps within the assembly with default parameters. Finally, we obtained an assembly of 638.70 Mb in size containing 5,490 scaffolds. The N50 values of contigs and scaffolds were 294.41 Kb and 2.61 Mb (Table 1), respectively, revealing a good contiguity of the genome assembly. The longest scaffold and contig were 9.02 Mb and 1.96 Mb, respectively. The assembled length was account for ~90 % of estimated genome size. To generate a chromosomal-level genome assembly, 133.17 Gb high-quality Hi-C data was used to further assembly. We first used HiC-Pro software (version2.8.0_devel) with default parameters to get ~26 Gb valid sequencing data, accounting for 19.31% of total Hi-C clean reads. Then, Juicer (version 1.5, an opensource tool for analyzing Hi-C datasets) ^12^, and the 3D de novo assembly pipeline ^13^ were used to connect the scaffolds to chromosomes with length of 603Mb (account for 94.40% of total genome) and scaffold sequences were assigned to 24 chromosomes, with the length from 14.13 Mb to 33.19 Mb (Table 2, Figure 2A and Figure 3).

**Figure 2.**
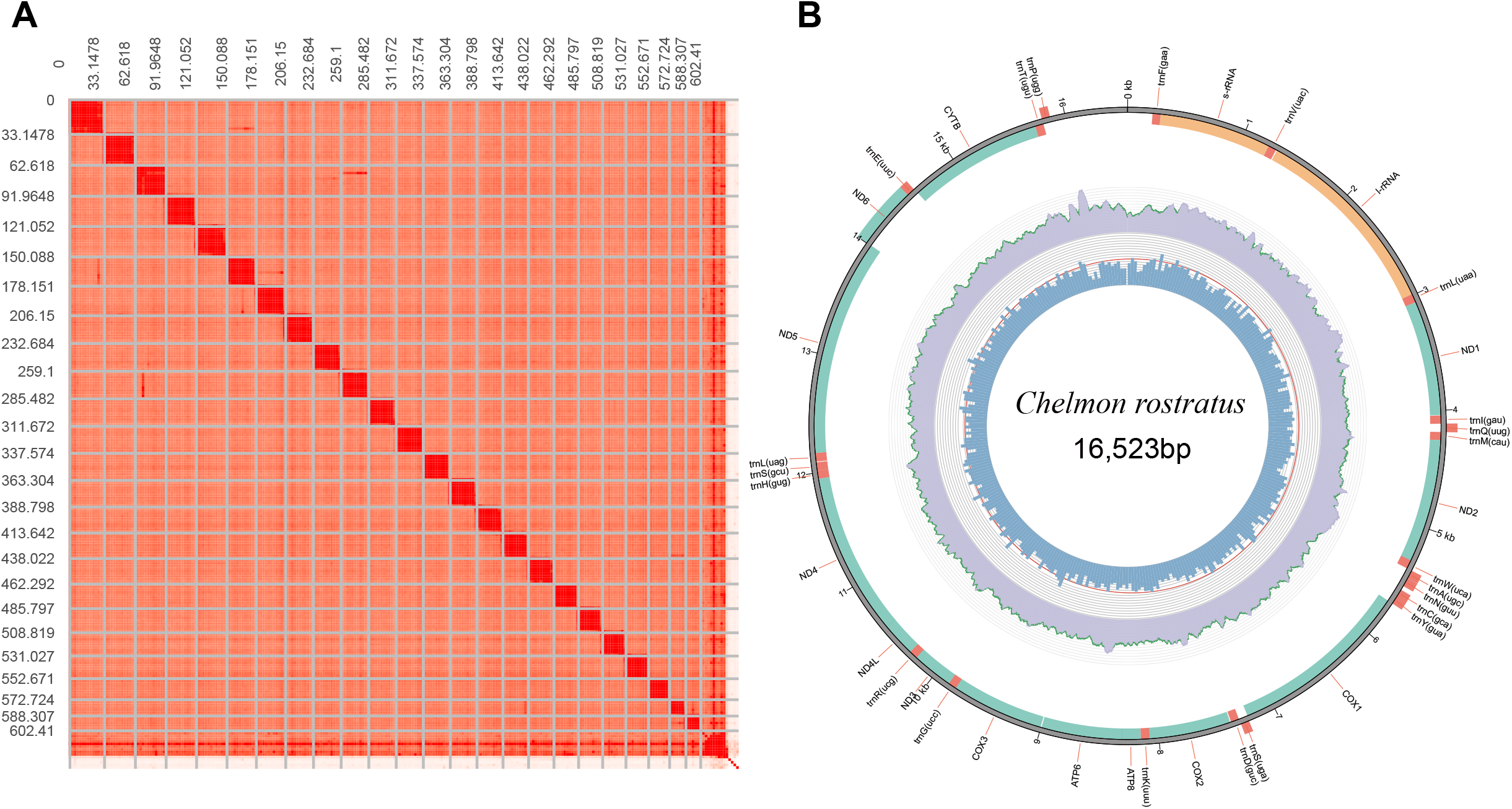
Heat map of interactive intensity between chromosome sequences (A) and Physical map of mitochondrial assembly (B).

**Figure 3.**
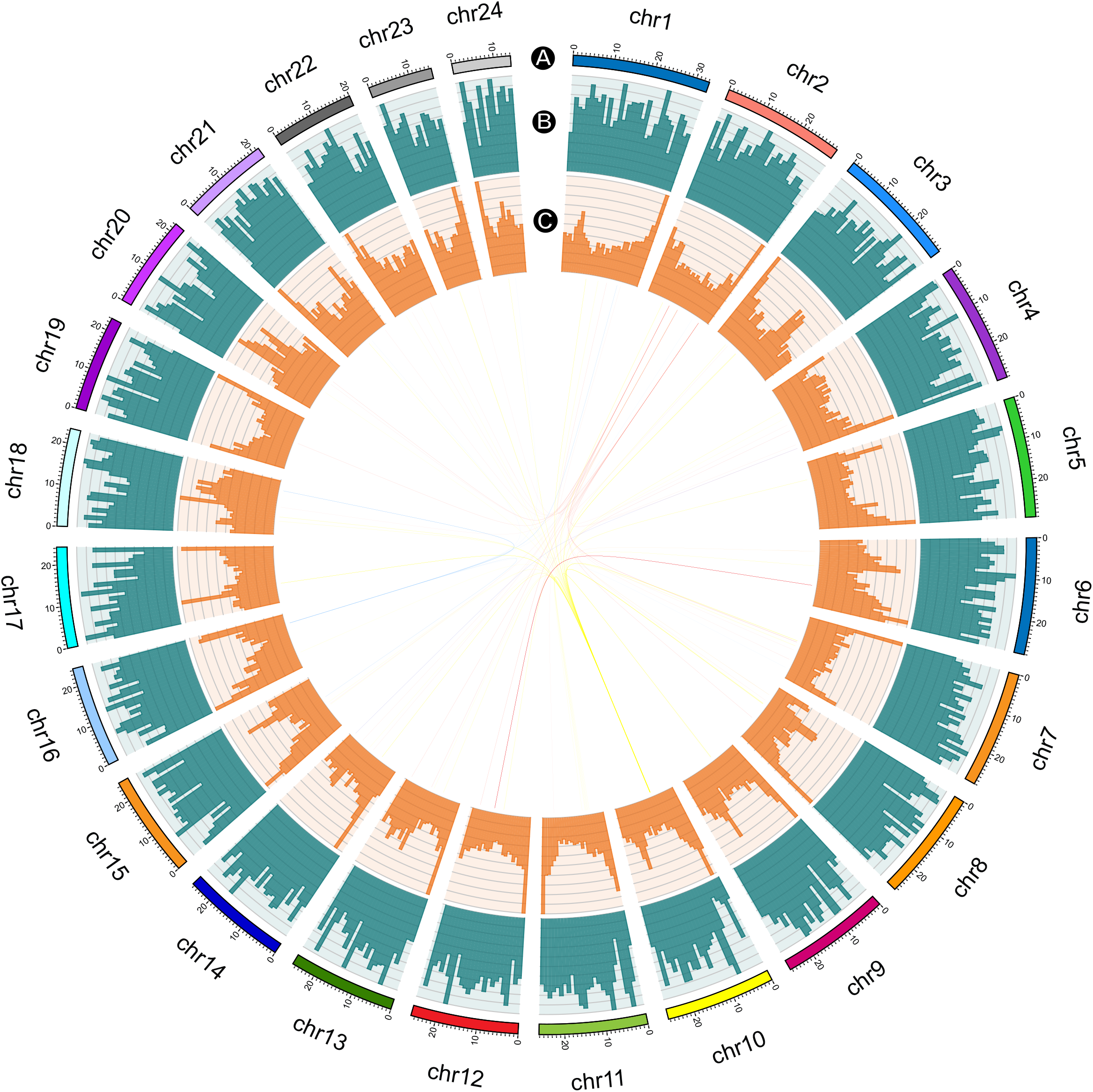
Distribution of basic genomic elements of *Chelmon rostratus* genome. (A) Chromosome karyotype. Different colored represented different chromosome we assembled. (B) Gene density. The histogram indicated number of genes per 1 Mb ranges from a minimum of 0.17 to a maximum of 1, illustrated by blue bar. (C) Repeat sequence density. The histogram indicated average DNA TE ratio per 1 Mb ranges from 0.16 to 1, illustrated by orange bar. Synteny blocks of each chromosome was illustrated by color lines, indicating that whole-genome duplication (WGD) event was not detected in *Chelmon rostratus* genome. Circos (Krzywinskiet al. 2009) (http://circos.ca) was used for constructing this diagram.

**Table 1.** Summary of the scaffold-level assembly. The N50 values of contigs and scaffolds were 294.41 Kb and 2.61 Mb, respectively, indicating good contiguity of the scaffold-level assembly for further Hi-C data assembly.

| Species name | *Chelmon rostratus* |
| --- | --- |
| Estimated genome size (bp) | 711,393,276 |
| Assembly size (bp) | 638,700,992 |
| Scaffold N50 (bp) | 2,610,810 |
| Longest scaffold (bp) | 9,024,273 |
| Contig N50 (bp) | 294,414 |
| Longest contig (bp) | 1,961,159 |
| Number of genes | 21,375 |
| GC content (%) | 42.52 |
| BUSCO (%) | 98.40 |
| Mapping ratio (%) | 90.76 |
| Coverage (%) | 99.28 |

**Table 2.** Summary of the chromosome-level assembly. About ~26 Gb valid Hi-C sequencing data was used to connect scaffold-level assembly into chromosomes with length of 603Mb (account for 94.40% of scaffold-level assembly size) and constructed 24 chromosomes, with lengths ranging from 14.13 (chr24) Mb to 33.19 Mb (chr1).

| Chromosome | Length (bp) | GC (%) |
| --- | --- | --- |
| chr1 | 33,188,845 | 41.94 |
| chr2 | 29,501,685 | 42.26 |
| chr3 | 29,388,801 | 42.61 |
| chr4 | 29,119,845 | 42.33 |
| chr5 | 29,062,128 | 42.38 |
| chr6 | 28,079,113 | 42.55 |
| chr7 | 28,043,518 | 42.42 |
| chr8 | 26,564,978 | 42.56 |
| chr9 | 26,447,630 | 42.51 |
| chr10 | 26,405,724 | 42.19 |
| chr11 | 26,164,899 | 42.37 |
| chr12 | 25,929,763 | 42.31 |
| chr13 | 25,766,544 | 42.17 |
| chr14 | 25,529,937 | 42.41 |
| chr15 | 24,867,447 | 42.81 |
| chr16 | 24,414,281 | 43.00 |
| chr17 | 24,306,480 | 42.65 |
| chr18 | 23,537,532 | 42.75 |
| chr19 | 23,051,343 | 42.21 |
| chr20 | 22,222,359 | 42.56 |
| chr21 | 21,649,575 | 42.91 |
| chr22 | 20,076,678 | 43.05 |
| chr23 | 15,560,595 | 43.28 |
| chr24 | 14,125,005 | 43.64 |
| Average | 25,125,196 | 42.58 |

We randomly selected 4 Gb clean data to assemble mitochondrial genome sequence using MitoZ software ^14^, resulting 16.52 kb of assembly with a cyclic structure (Figure 2B).

### Repetitive sequence and gene annotation

We annotated the two major types of repetitive sequences (tandem repeats, TRFs and transposable elements, TEs) in the assembled genome. According to the method of previously research ^15–17^, TRFs were identified using Tandem Repeats Finder (version 4.04) ^18^. Transposable elements (TEs) were identified by a combination of homology-based and *de novo* approaches. Briefly, for homology-based annotation, known repeats in the database (Repbase16.02) ^19^ were aligned against the genome assembly using RepeatMasker and RepeatProteinMask (version 3.2.9) ^20^ at both the DNA and protein levels. For *de novo* annotation, RepeatModeler (version 1.1.0.4) ^21^ was employed to build a *de novo* non-redundant repeat library and then this repeat library was searched against the genome using RepeatMasker ^20^. In this way, up to 21.47 % of the assembled sequences were found to be repeat sequences (Figure 3, Supplementary Table 3).

Protein-coding gene were then predicted by a combination of two ways: (1) the *ab initio* gene prediction and (2) the homology-based annotation ^22–24^. For *ab initio* gene prediction approaches, Augustus ^25^ and GlimmerHMM ^26^ were used with *Danio rerio* as the species of HMM model to predict gene models; For homology-based annotation, four homolog species including *Pundamilia nyererei, Maylandia zebra, Astatotilapia calliptera* and *Perca flavescens* were aligned against the genome assembly using BLAT software (version 0.36) ^27^ and GeneWise software (version 2.4.1) ^28^. 21,375 protein-coding genes were obtained by combining the different evidences using Glean software (version 1.0) ^29^. In the final gene models, the average length was 16,183.81 bp, with an average of 10 exons. The average length of coding sequences, exons and introns were 1,789.82bp, 179.10 bp and 1599.48 bp, respectively, similar to that of the other released fish genomes, such as *Astatotilapia calliptera, Maylandia zebra, Perca flavescens* and *Pundamilia nyererei* ^30–32^ (Supplementary Table 4, Supplementary Figure 2). Gene annotation of mitochondria was performed using MitoZ software ^14^, and 13 protein-coding genes as well as 22 tRNA genes were annotated.

Functions of the annotated protein-coding genes were inferred by searching homologs in the databases (KEGG, COG, NR, Swissprot and Interpro) ^33–37^. In this way, 18,005 (84.23%), 7,343 (34.35%), 20,141 (94.23%), 19,114 (89.42%) and 19,313 (90.35%) of protein-coding genes had their homologous alignment in the above databases, respectively. The remaining 1,212 (5.67%) protein-coding genes with unknown function might be the specific feature of the *Chelmon rostratus* genome (Supplementary Table 5).

### Gene family and phylogenetic analysis

To identify and analyze the gene families, we selected other eleven species with whole genome sequences available *(Lepisosteus oculatus, Hippocampus comes, Larimichthys crocea, Gasterosteus aculeatus, Takifugu rubripes, Oreochromis niloticus, Astyanax mexicanus, Danio rerio, Cynoglossus semilaevis, Oryzias latipes* and *Homo sapiens* as outgroup) ^24,38–47^. The protein coding genes of the total twelve species were clustered into 18,502 gene families using TreeFam ^48–50^. Among these gene families, 2,301 were single-copy gene families (one copy in each of these species) (Figure 4B). The 21,375 protein-coding genes of *Chelmon rostratus* were classified into 13,797 gene families, given an average of 1.55 genes per gene family. Compared to the other species, it was similar to the gene family numbers of *Lepisosteus oculatus* (13,967) and *Gasterosteus aculeatus* (13,331), but was quite different from those of *Larimichthys crocea* (14,724), *Homo sapiens* (14,578) and *Takifugu rubripes* (12,554) (Supplementary Table 6). Among the clustered gene families from *Chelmon rostratus*, 8,190 gene families were common to at least one of the other species, the remaining 52 gene families were unique. Between the four species *(Chelmon rostratus, Danio rerio, Takifugu rubripes* and *Larimichthys crocea)*, the number of common shared and unique gene was shown in Figure 4C. To understand the function of these gene families, we further performed GO enrichment with these gene families from *Chelmon rostratus*, compared with the other 11 species. The result reflected that unique gene families from *Chelmon rostratus* were enriched in muscle contraction functions (Supplementary Table 7). Phylogenetic analysis using the concatenated sequence alignment of the 2,301 single-copy genes shared by the twelve species was performed. The PhyML software (version 3.0) ^51^, based on the method of maximum likelihood, was used to construct the phylogenetic tree. The split time between *Chelmon rostratus* and *Larimichthys crocea* was estimated to be ~92 million years ago (Figure 4A). Based on the similarity of the protein sequences, 483 syntenic blocks were identified by using McScanX software (version 0.8) ^52^ (Supplementary Table 8). The time of the duplication and divergence event in these species was calculated based on the distribution of synonymous mutation rate for the gene pairs in the paralogous syntenic blocks, indicating that whole-genome duplication (WGD) event was not detected in *Chelmon rostratus* genome (Figure 3). The expansion and contraction of the gene family analysis may reveal the evolutionary dynamics of gene families thus provide the clues for understanding the diversity of different species. It is often inferred from the number of genes in the gene family and the phylogenetic tree. In our study, we used the CAFÉ (version 2.1) ^53^ software to analyze the expansion and contraction of clustered gene families (Figure 4D). As a result, a total of 18,498 gene families from the most recent common ancestor (MRCA) have been identified. Compared to the recent common ancestor between *Chelmon rostratus* and *Larimichthys crocea*, 793 gene families were expanded and the majority of the expanded gene families were found to be involved in synapse organization. (Supplementary Table 9). On the other hand, there was 2,962 gene family contracted involved in immune system process (Supplementary Table 10).

**Figure 4.**
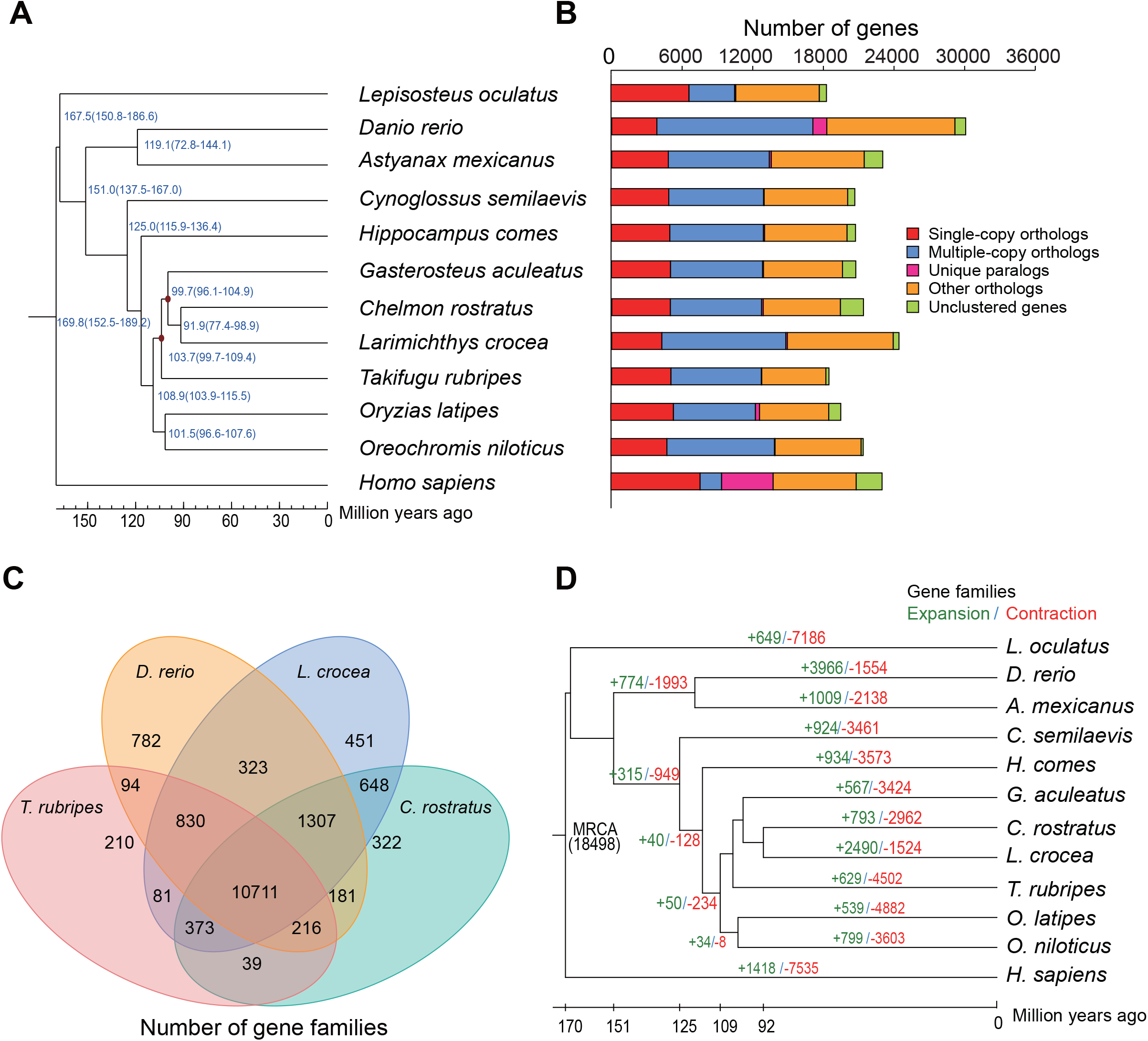
Comparative analysis of the *Chelmon rostratus* genome. (A) Phylogenetic analysis among *Lepisosteus oculatus, Hippocampus comes, Larimichthys crocea, Gasterosteus aculeatus, Takifugu rubripes, Oreochromis niloticus, Astyanax mexicanus, Danio rerio, Cynoglossus semilaevis; Oryzias latipes* and *Homo sapiens* as outgroup, by using the single-copy gene families. The species differentiation time between *Chelmon rostratus* and *Larimichthys crocea* was ~92 million years ago. (B) The protein coding genes of the total twelve species were clustered into 18,502 gene families. Among these gene families, 2,301 were single-copy gene families (one copy in each of these species). (C) Venn diagram showing overlaps of gene families between *Chelmon rostratus, Danio rerio, Takifugu rubripes* and *Larimichthys crocea*. A total of 322 gene families were unique to *Chelmon rostratus* and 10,711 were commonly shared by the other species genome. (D) Compared to the recent common ancestor between *Chelmon rostratus* and *Larimichthys crocea*, 793 gene families were expanded and 2,962 gene family contracted in *Chelmon rostratus* genome.

### Data Records

Raw reads from BGISEQ-500 sequencing are deposited in the CNGB Nucleotide Sequence Archive (CNSA) with accession number CNP0000597 (https://db.cngb.org/cnsa). Data Citation 1: CNGB Nucleotide Sequence Archive CNP0000597).

### Technical Validation

To evaluate the genome assembly, we aligned sequencing data which we filtered previously using SOAPaligner (version 2.2) ^10^ and found that 90.76% could be mapped back to the assembled genome. we also calculate its GC depth to rule out possible biases during sequencing or possible contaminations. We identified the average GC contents of this genome to be ~42.52% and we found a continuous GC depth distribution (Supplementary Figure 3), indicating no obvious assembly errors resulted from GC content or contamination. The genome completeness assessment was estimated with Benchmarking Universal Single-copy Orthologs (BUSCO, version 3.0.1) ^54^. BUSCO analysis showed that 97.3% (2,518) of core genes were found in our assembly with 2,491 (96.3%) were single copy gene and 27 (1.0%) were duplicated (Supplementary Table 11), indicating a good coverage of the genome.

To validate the quality of predicted gene sets, we also assessed the completeness using BUSCO (version 3) with the fish core gene database (actinopterygii_odb9) ^55^. We found that about 90.20% of core gene were annotated in our gene set with 4,000 were single copy gene and 132 were duplicated (Supplementary Table 12).

## Acknowledgements

This work is supported by the special funding of “Blue granary” scientific and technological innovation of China (2018YFD0900301-05). The work also received the technical support from China National Gene Bank.

## Author contributions

Y.S., G.F., S.L. and X.L. conceived the work. M.Z. collected sample and X.H. sequenced the libraries. Y.S., X.H., S.Z., Y.A. collected the public data and performed the analyses. Y.C., M.L., C.L. and Y.Z. helped in the analysis. Y.S., X.H., G.F., I.S., S.L. and X.L. wrote and revised the manuscript.

