## Supplemental Tables for "Chromosome-level genome assembly of a butterflyfish, *Chelmon rostratus*"

**Supplementary Table 1** Summary of the library types and data generated in this study.

|  | Raw data | | Clean data | |
| --- | --- | --- | --- | --- |
| Library type | Total data  (bp) | Sequence  depth (X) | Total data  (bp) | Sequence  depth (X) |
| stLFR | 134,636,786,200 | 210.74 | 59,539,880,400 | 93.19 |
| Hi-C | 134,180,382,000 | 210.02 | 133,174,561,400 | 208.45 |

**Supplementary Table 2** *k*-mer analysis and statistics of estimated genome size.

|  | *k*-mer | *k*-mer num | Peak depth | Estimated genome size | Used base |
| --- | --- | --- | --- | --- | --- |
|  | 17 | 44,106,383,087 | 62 | 711,393,276 | 53,260,871,600 |

**Supplementary Table 3**	Statistics of repetitive sequence annotation.

|  | Type | Percentage (%) | Length  (bp) |
| --- | --- | --- | --- |
| **Type I: Retrotransposon elements** | |  |  |
|  | SINE | 0.08 | 574,726 |
|  | LINE | 4.15 | 26,502,977 |
|  | LTR | 3.22 | 20,579,971 |
|  | Other | 0.00 | 0 |
| **Type II: DNA transposon** | |  |  |
|  | DNA | 7.91 | 50,536,086 |
| **Type III: Tandem repeats** | |  |  |
|  | Satellite | 0.10 | 690,841 |
|  | Simple repeat | 1.17 | 7,517,943 |
| **unknow** |  | 5.75 | 36,756,742 |
| **Total repeat** |  | 21.47 | 137,130,362 |

**Supplementary Table 4**	Statistics of gene model in different species.

| Species | # of genes | | Average gene length (bp) | Average CDS length (bp) | Average exon number | Average exon length (bp) | Average intron length (bp) |
| --- | --- | --- | --- | --- | --- | --- | --- |
| *Astatotilapia calliptera* | | 50,280 | 20,912.17 | 2,312.05 | 12.85 | 179.90 | 1,569.39 |
| *Maylandia zebra* | | 14,016 | 18,837.56 | 2,191.49 | 12.65 | 173.23 | 1,428.78 |
| *Perca flavescens* | | 42,830 | 16,993.63 | 2,001.65 | 11.68 | 171.43 | 1,404.25 |
| *Pundamilia nyererei* | | 38,343 | 18,382.06 | 2,045.40 | 11.74 | 174.29 | 1,521.78 |

**Supplementary Table 5** Summary of homologous function annotation through several databases.

| Database | Number | Percentage (%) |
| --- | --- | --- |
| Nr | 20,141 | 94.23 |
| Swissprot | 19,114 | 89.42 |
| KEGG | 18,005 | 84.23 |
| COG | 7,343 | 34.35 |
| Interpro | 19,313 | 90.35 |
| Total | 20,163 | 94.33 |

**Supplementary Table 6**	Statistics of gene families.

| Species | Genes number | Genes in families | Unclustered genes | Family number | Unique families | Average genes per family |
| --- | --- | --- | --- | --- | --- | --- |
| *L. oculatus* | 18,313 | 17,707 | 606 | 13,967 | 38 | 1.27 |
| *H. comes* | 20,732 | 20,017 | 715 | 13,975 | 46 | 1.43 |
| *L. crocea* | 24,403 | 23,914 | 489 | 14,724 | 46 | 1.62 |
| *G. aculeatus* | 20,756 | 19,626 | 1,130 | 13,331 | 20 | 1.47 |
| *T. rubripes* | 18,459 | 18,200 | 259 | 12,554 | 11 | 1.45 |
| *O. niloticus* | 21,431 | 21,252 | 179 | 13,306 | 14 | 1.6 |
| *A. mexicanus* | 23,024 | 21,455 | 1,569 | 14,209 | 73 | 1.51 |
| *D. rerio* | 30,067 | 29,157 | 910 | 14,444 | 140 | 2.02 |
| *C. semilaevis* | 20,722 | 20,118 | 604 | 14,130 | 22 | 1.42 |
| *O. latipes* | 19,535 | 18,515 | 1,020 | 12,815 | 82 | 1.44 |
| *H. sapiens* | 23,032 | 20,838 | 2,194 | 14,578 | 647 | 1.43 |
| *C. rostratus* | 21,375 | 19,421 | 1,954 | 13,797 | 52 | 1.41 |

**Supplementary Table 7** GO enrichment analysis of 52 unique gene families in *Chelmon rostratus* genome.

| GO terms | Description | q-value |
| --- | --- | --- |
| GO:0016820 | hydrolase activity, acting on acid anhydrides, catalyzing transmembrane movement of substances | 2.66E-04 |
| GO:0035556 | intracellular signal transduction | 2.41E-03 |
| GO:0017022 | myosin binding | 4.22E-08 |
| GO:0006915 | apoptotic process | 4.62E-03 |
| GO:0003779 | actin binding | 2.26E-03 |
| GO:0007205 | protein kinase C-activating G-protein coupled receptor signaling pathway | 4.05E-04 |
| GO:0005516 | calmodulin binding | 7.58E-05 |
| GO:0004143 | diacylglycerol kinase activity | 4.05E-04 |
| GO:0003723 | RNA binding | 4.80E-02 |
| GO:0003333 | amino acid transmembrane transport | 3.57E-05 |
| GO:0016491 | oxidoreductase activity | 1.75E-02 |
| GO:0055114 | oxidation-reduction process | 4.67E-02 |
| GO:0006811 | ion transport | 2.07E-02 |
| GO:0008234 | cysteine-type peptidase activity | 3.28E-03 |
| GO:0005524 | ATP binding | 2.31E-02 |
| GO:0004652 | polynucleotide adenylyltransferase activity | 4.22E-08 |
| GO:0015991 | ATP hydrolysis coupled proton transport | 2.38E-03 |
| GO:0033178 | proton-transporting two-sector ATPase complex, catalytic domain | 2.66E-04 |
| GO:0004197 | cysteine-type endopeptidase activity | 8.62E-04 |
| GO:0005861 | troponin complex | 3.30E-05 |
| GO:0043631 | RNA polyadenylation | 4.22E-08 |
| GO:0006936 | muscle contraction | 1.27E-07 |
| GO:0015171 | amino acid transmembrane transporter activity | 3.57E-05 |

**Supplementary Table 8**	Statistics of synteny blocks.

| Syntenic Blocks | Average Syntenic Gene Pairs Per Block | Syntenic Gene Pairs | Mean Block Length |
| --- | --- | --- | --- |
| 483 | 8.1656 | 3,944 | 1,656,180.17 |

**Supplementary Table 9** GO enrichment analysis of 793 expanded gene families in *Chelmon rostratus* genome.

| GO terms | Description | q-value |
| --- | --- | --- |
| GO:0042043 | neurexin family protein binding | 4.64E-03 |
| GO:0005856 | cytoskeleton | 2.55E-02 |
| GO:0016021 | integral component of membrane | 1.09E-09 |
| GO:0019069 | viral capsid assembly | 0.00E+00 |
| GO:0005891 | voltage-gated calcium channel complex | 6.95E-03 |
| GO:0007186 | G-protein coupled receptor signaling pathway | 0.00E+00 |
| GO:0060076 | excitatory synapse | 2.26E-04 |
| GO:0008146 | sulfotransferase activity | 1.62E-14 |
| GO:0002682 | regulation of immune system process | 0.00E+00 |
| GO:0005245 | voltage-gated calcium channel activity | 6.07E-03 |
| GO:0006334 | nucleosome assembly | 2.51E-02 |
| GO:0004252 | serine-type endopeptidase activity | 1.01E-02 |
| GO:0004984 | olfactory receptor activity | 0.00E+00 |
| GO:0050804 | modulation of synaptic transmission | 1.92E-03 |
| GO:0070588 | calcium ion transmembrane transport | 2.41E-02 |
| GO:0004930 | G-protein coupled receptor activity | 0.00E+00 |
| GO:0050808 | synapse organization | 1.24E-03 |

**Supplementary Table 10** GO enrichment analysis of 2,962 contracted gene families in *Chelmon rostratus* genome.

| GO terms | Description | q-value |
| --- | --- | --- |
| GO:0042043 | neurexin family protein binding | 4.64E-03 |
| GO:0001594 | trace-amine receptor activity | 0.00E+00 |
| GO:0016021 | integral component of membrane | 1.09E-09 |
| GO:0006955 | immune response | 1.02E-04 |
| GO:0042605 | peptide antigen binding | 1.26E-07 |
| GO:0007186 | G-protein coupled receptor signaling pathway | 0.00E+00 |
| GO:0002474 | antigen processing and presentation of peptide antigen via MHC class I | 1.26E-07 |
| GO:0004252 | serine-type endopeptidase activity | 1.01E-02 |
| GO:0004984 | olfactory receptor activity | 0.00E+00 |
| GO:0042613 | MHC class II protein complex | 4.64E-03 |
| GO:0042612 | MHC class I protein complex | 1.26E-07 |
| GO:0019882 | antigen processing and presentation | 6.26E-03 |
| GO:0005622 | intracellular | 4.82E-03 |
| GO:0004930 | G-protein coupled receptor activity | 0.00E+00 |

**Supplementary Table 11**	Evaluation of chromosome-level assembly.

| Item | Quantity | Proportion (%) |
| --- | --- | --- |
| Complete BUSCOs | 2,518 | 97.3 |
| Complete and single-copy BUSCOs | 2,491 | 96.3 |
| Complete and duplicated BUSCOs | 27 | 1.0 |
| Fragmented BUSCOs | 48 | 1.9 |
| Missing BUSCOs | 20 | 0.8 |
| Total BUSCO groups searched | 2,586 | NA |

**Supplementary Table 12**	Evaluation of predicted gene models.

| Type | Number | Percentage (%) |
| --- | --- | --- |
| Complete BUSCOs | 4,132 | 90.2 |
| Complete and single-copy BUSCOs | 4,000 | 87.3 |
| Complete and duplicated BUSCOs | 132 | 2.9 |
| Fragmented BUSCOs | 288 | 6.3 |
| Missing BUSCOs | 164 | 3.5 |
| Total BUSCO groups searched | 4,584 | NA |
