## Supplementary figures and images for "Chromosome-level genome assembly of a butterflyfish, *Chelmon rostratus*"

### FigureS1

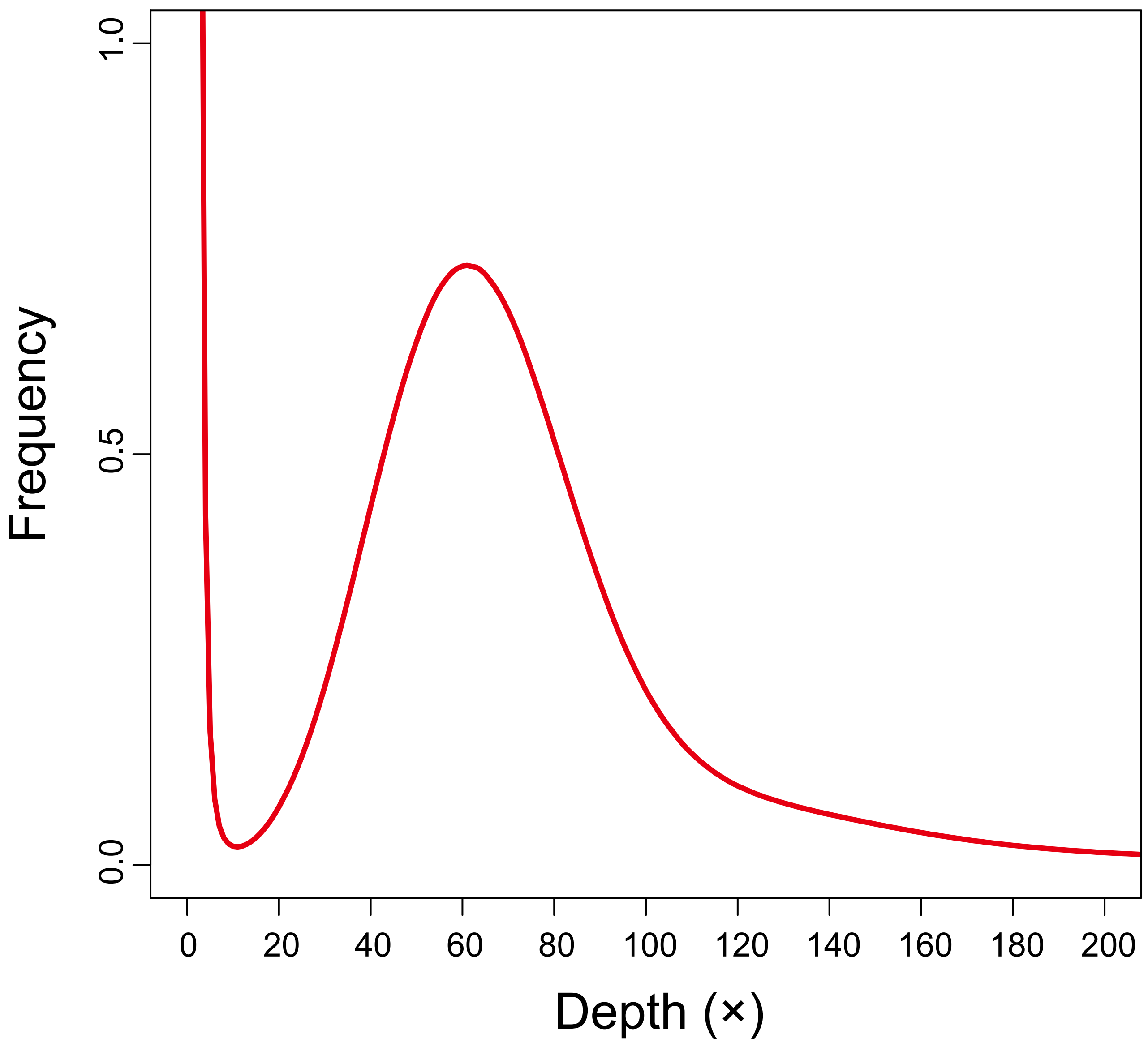

### FigureS2

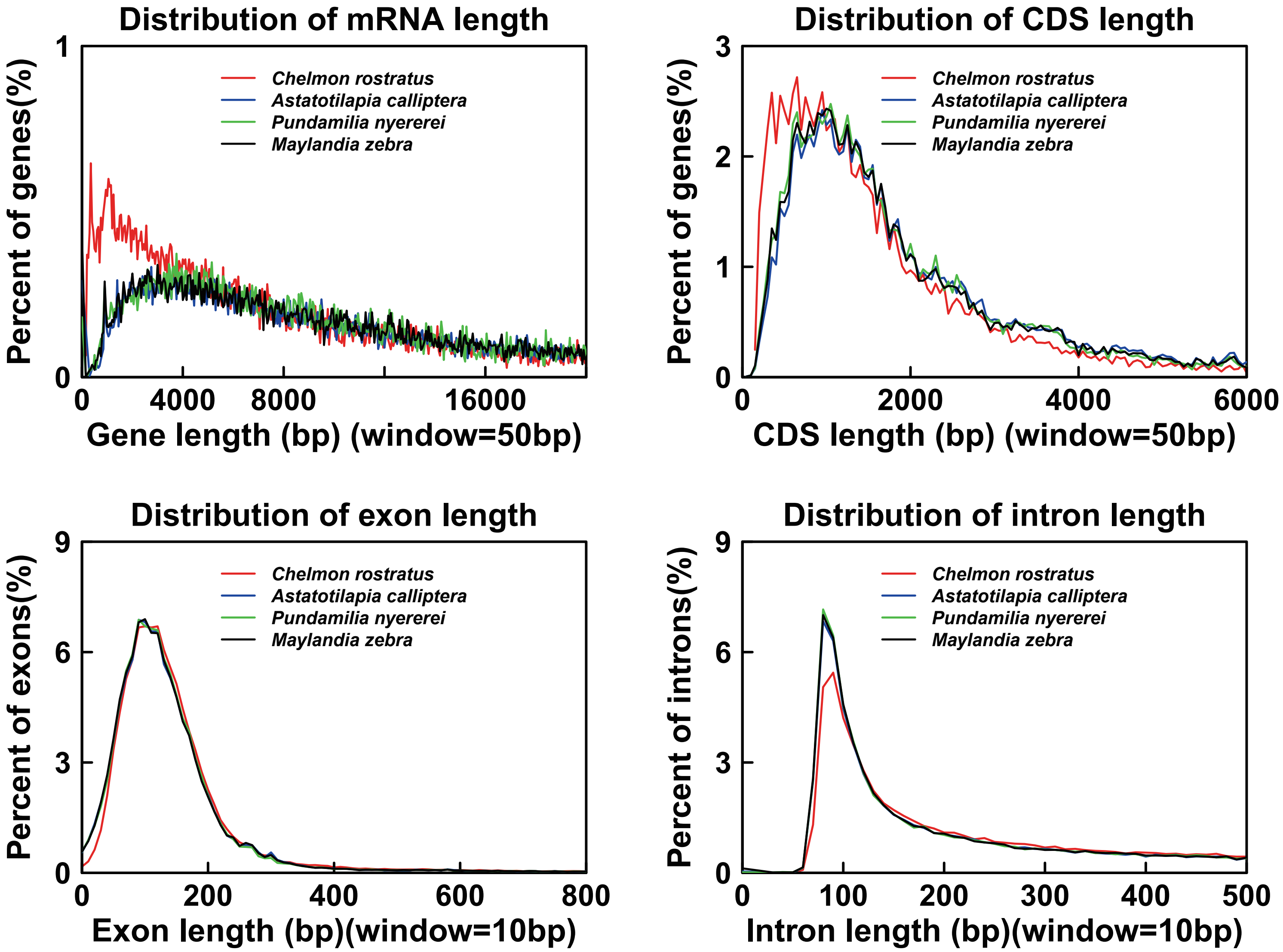

### FigureS3

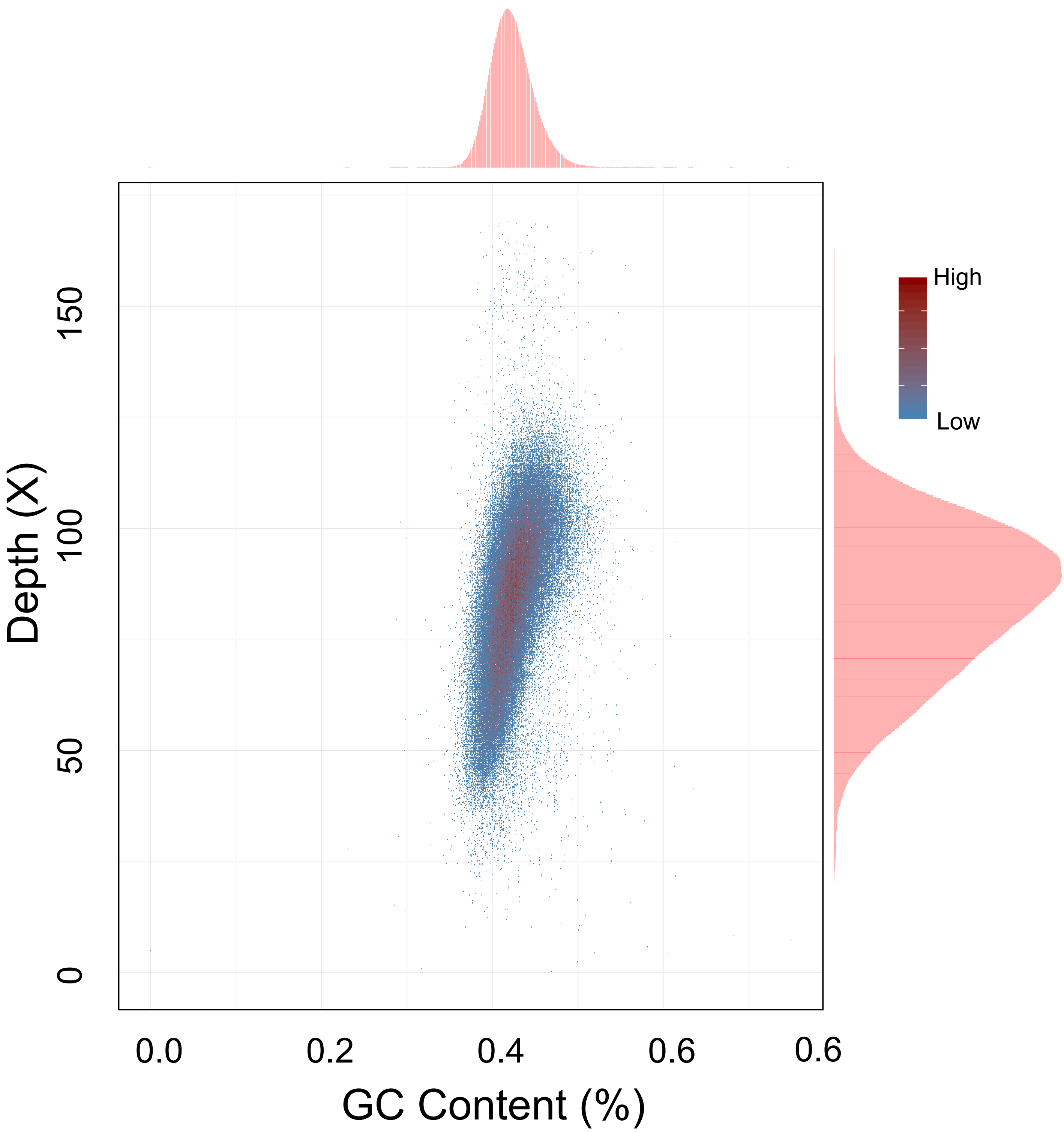
